# Phenotypic divergence associated with genomic changes suggest local adaptation in obligate asexuals

**DOI:** 10.1101/2024.08.15.608098

**Authors:** Athina Karapli-Petritsopoulou, Jasmin Josephine Heckelmann, N. John Anderson, Sören Franzenburg, Dagmar Frisch

## Abstract

Local adaptation is a key evolutionary process that generates global biodiversity and is promoted by environmental heterogeneity. Most of the existing knowledge on adaptive capacity is focused on sexual organisms, while the adaptation potential of asexually reproducing eukaryotes as well as their vulnerability to environmental change remains unclear. Cyclical parthenogens of the keystone freshwater grazer *Daphnia* are known to be locally adapted with genetic differentiation related to environmental conditions even between neighbouring ponds. Similar patterns have been found in obligate parthenogenetic congeners, however studies on their local adaptation potential are rare. Here, we use respiration rate and whole genome sequencing to test the local adaptation of Arctic asexual polyploid *Daphnia* from lakes with contrasting oxygen environments. The genomic data revealed the presence of two genetic clusters which differed significantly in respiration rates, suggesting molecular evolution as an adaptive mechanism. Functional enrichment pointed to differences in metabolic processes between the two genetic clusters. Our results, combining phenotypic and whole genome sequencing data, suggest that these clones are locally adapted to low oxygen concentrations and provide support for local adaptation by evolution in an obligate parthenogen.

## Introduction

Environmental heterogeneity creates opportunities for local adaptation to evolve, thereby enhancing biodiversity. Locally adapted resident genotypes have a higher habitat-specific mean fitness than genotypes from a different habitat (Kawecki and Ebert 2004), potentially giving rise to subsequent speciation (Schluter 2001). At the current unprecedented rate of global environmental change, local adaptation can shift to maladaptation and prove to be a cost (Derry et al. 2019; Anderson and Wadgymar 2020). Investigating the extent of local adaptation and the mechanisms for its evolution in contrasting environments can help to understand the potential for evolutionary rescue (Schiffers et al. 2013) and to guide informed conservation measures for the protection of biodiversity in a changing world (Meek et al. 2023).

Asexually reproducing eukaryotes might be less common than sexuals but are spread throughout the tree of life in multiple independently evolved lineages (Lodé 2013). They are known to mostly occupy “marginal” and disturbed habitats in comparison to their sexual counterparts, a phenomenon known as geographic parthenogenesis (Tilquin and Kokko 2016). Asexual reproduction offers an advantage for rapid colonization and propagation in new habitats (Hörandl et al. 2008). In the long-term, asexual lineages are generally considered evolutionary dead ends and short-lived due to a lack of genetic variation and the occurrence of mutational meltdown through the accumulation of deleterious mutations (Felsenstein 1974; Lynch et al. 1993). Ancient asexual groups such as bdelloid rotifers, and darwinulid ostracods are a counterexample for these arguments (Schwander and Crespi 2009), although recent studies have uncovered cryptic sex in the first group (Laine et al. 2022). Despite the long-standing interest in the enigma of sex, the adaptation potential of asexually reproducing populations to contrasting habitats or their ability to cope with rapid environmental change is still unclear. More whole genome studies are needed to shed added light on the genomic evolution of asexuals (Hörandl et al. 2020).

Asexuality in eukaryotes is often coupled with polyploidy, which could explain the ecological advantage of asexual polyploids better than their mode of reproduction (Lundmark and Saura 2006). Polyploidy can have complex effects on the adaptation potential of an organism, making it difficult to formulate a general rule based on findings from different species (Van de Peer et al. 2017). The larger cell and body size that comes with polyploidy has been suggested as a main reason for the increased abiotic tolerance of polyploids as it affects many biological processes (Doyle and Coate 2019). Polyploidy can also offer an advantage due to heterosis in cases of allopolyploids or gene redundancy, which can mask deleterious recessive alleles or reduce the constraint on neo- or sub-functionalization of gene copies allowing more opportunities for adaptive functions (Comai 2005). Moreover, polyploid populations have a higher chance of novel beneficial mutations arising and possibly a faster adaptation rate (Otto and Whitton 2000). The above characteristics could ameliorate the evolutionary fate of asexual polyploids.

The cladoceran *Daphnia*, a keystone grazer in freshwater systems, is well-known for its ability to adapt rapidly to environmental change, e.g. as shown in studies of eutrophication (Hairston et al. 1999; Frisch et al. 2014) and predation pressure (Cousyn et al. 2001). Commonly, it reproduces by cyclical parthenogenesis (CP). This mode of reproduction combines female asexual propagation and sexual reproduction to produce dormant eggs encased in ephippia towards the end of the season (Miner et al. 2012). The alteration between the two modes of reproduction seems to be most beneficial for rapid local adaptation since it retains the benefit of rapid propagation followed by clonal selection and can generate genetic variation through recombination during sexual reproduction (Lynch and Gabriel 1983; De Meester et al. 2016). After a dormant period, the resting eggs continue development when conditions are favourable and function both as a dispersal agent for colonization of new habitats (Incagnone et al. 2015), and as members of egg banks for local recruitment of the *Daphnia* population in the next season (Hairston Jr. 1996). These often very large egg banks can provide a buffer against possibly maladapted new colonizers entering local populations and thus safeguard local adaptations from gene flow (De Meester et al. 2002). Indeed, in studies of ponds located very close to each other but with contrasting environmental conditions, *Daphnia* populations were found to be genetically divergent (Vanoverbeke and De Meester 1997) and locally adapted to environmental factors such as differing oxygen concentration (Carvalho 1984), pond water (Declerck et al. 2001), and fish predation pressure (De Meester 1996b). While CP lines show this consistent pattern of local adaptation, work on obligate parthenogenetic (OP) species is less conclusive.

The *Daphnia pulex* species complex includes various OP lineages that inhabit higher altitudes and latitudes and are often polyploid (Decaestecker et al 2009). OP *Daphnia* reproduce asexually both during the growing season and for ephippia production. Additionally, some OP lineages of the *Daphnia pulex* complex are able to produce functional males, which when mated with CP females may cause asexuality either through contagious asexuality or as a byproduct of hybridization (Innes and Hebert 1988; Xu et al. 2015). OP *Daphnia* are successful colonizers and can outcompete their sexual congeners after invasion as seen in some African lakes (Mergeay et al. 2006).

Studies of population genetic differentiation from both ponds and lakes inhabited by OP *Daphnia* have shown the same pattern as in cyclical parthenogens: genetic differentiation matches environmental variability. More specifically, microgeographic adaptation to a salinity gradient among neighbouring rock pools in Canada was found in OP lineages of the Arctic *Daphnia pulex/pulicaria* species complex (Weider and Hebert 1987; Weider et al. 2010), while patterns of environmental sorting to food availability and predation risk were found in OP *Daphnia pulex* lineages from lakes in the Andes (Aguilera et al. 2007). Similarly, Haileselasie et al. (2016a) found environmental variables to be a main explanatory factor for the genetic composition of *Daphnia* OP lineages among ponds and lakes in the Jakobshavn Ibræ area in Greenland. These results, however, were restricted to low genomic resolution, and thus lack the association of habitat-specific physiological adaptations to genome differentiation. Finally, studies in Japanese OP *Daphnia pulex* lineages showed the occurrence of molecular evolution connected to heritable phenotypic traits, albeit dominated by few possibly pleiotropic mutations (So et al. 2015; Tian et al. 2019).

In this study, we present data from OP, polyploid Arctic *Daphnia pulicaria* clones originating from four neighbouring lakes in Southwest Greenland with contrasting environments. These four lakes historically (about 8000 y.a.) formed a single paleolake, and have separated since (Aebly and Fritz 2009). One of the most striking environmental aspects is the existence of a permanent anoxic zone in two of the four lakes. This anoxic zone is inhabited by purple sulfur bacteria, a potential food resource for the local *Daphnia* population (Dane et al., 2020). However, to be able to exploit this food source, *Daphnia* would have to feed in or near the oxycline which has strong implications on their ability to tolerate hypoxic conditions. Previous studies in several lakes in this area have shown low but lake-specific genetic diversity in the *Daphnia* populations (Dane et al, 2020).

Here, we measured respiration rates as an indicator of local adaptation that would allow tolerance to hypoxia in *Daphnia* originating from each of the four lakes, and performed whole genome sequencing to assess the genetic differentiation among the tested clonal lineages. Respiration rate is directly related to the aerobic metabolic rate which can be decreased by many taxa as part of a hypometabolic response to hypoxia, including *Daphnia* (Wiggins and Frappell 2000; Paul et al. 2004; Horscroft et al. 2017; Regan et al. 2017). We hypothesized that (1) *Daphnia* from the two lakes with an anoxic zone would be phenotypically adapted to hypoxia, hence have lower respiration rates, and (2) would be genetically more closely related to each other than to those of the two completely oxic lakes.

## Materials and Methods

### Study site

The study lakes are located at the head of Kangerlussuaq fjord in the arid interior of SW Greenland (Fig. 1). The climate is continental low arctic with a mean annual temperature of –6.2° C and annual precipitation ∼200 mm yr^−1^ although evaporation greatly exceeds precipitation during the summer months (JJA). The vegetation surrounding the lakes is dwarf shrub tundra with *Salix glauca, Betula nana*, *Ledum palustre spp. decumbens*, *Vaccinium uliginosum,* and *Rhododendron lapponicum* as co-dominants. Vegetation cover is variable due to water stress, with large areas of bare ground, in particular on south-facing slopes (see Anderson 2020 for further details). The lakes are located close to each other (∼5 km) and were all part of the large fossil lake system that formed after deglaciation (Anderson and Leng 2004; Aebly and Fritz 2009) (Fig. 1). They range in size from 60 ha to 300 ha and maximum water depths range from 12 to 30 m. Today, the lakes are all closed-basin systems and oligosaline (2400–5000 μS cm^−1^) due to evaporative concentration. SS5 and SS6 were separate lakes when first sampled in 1996 (Anderson et al 2001) but joined together as regional lake levels increased from 1999. Today they are separated by a shallow (1-m) sill and are functionally different limnologically. All four lakes thermally stratify during the ice-free period (Fig. 1b) and two (SS4, SS6) are meromictic (permanently, chemically stratified). SS3 and SS5 are large, deep basins and have oxygenated water columns (Fig. 1a) while SS4 is anoxic below 10 m and has a purple sulfur bacterial plate, observed during field sampling (XXX, unpublished field observations) and confirmed by abundant okenone in the sediments (McGowan et al. 2008). Okenone is the carotenoid associated with purple sulfur bacteria. SS6 is intermediate in terms of its O_2_ profile (see Fig. 1a) but again the presence of okenone and laminated sediments (McGowan et al. 2008) indicates that there is a small permanently anoxic sub-basin. All four lakes are fishless (Jeppesen et al. 2017) and contain asexual triploid *Daphnia pulicaria* populations (Dane et al. 2020). The oxygen and temperature depth profiles shown in Figure 1 were obtained during the sampling field campaign in August 2022 (for more details see the Supplementary material).

**Fig. 1.**
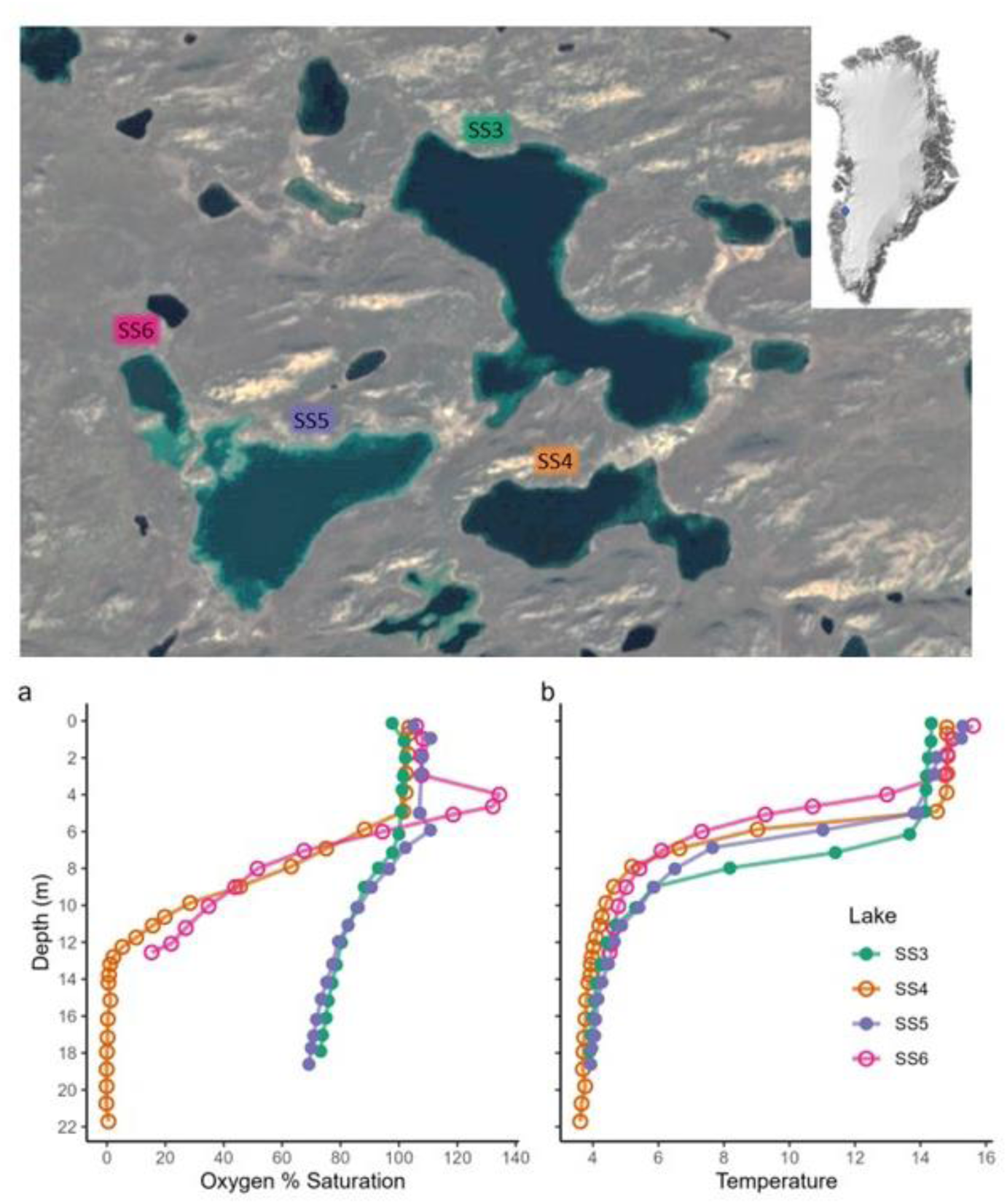
Top panel: Map of the four lakes used for this study in the Kangerlussuaq area, SW Greenland (formatted from Google maps). Bottom panel: **a.** Oxygen % Saturation depth profiles and **b.** Temperature depth profiles of the four lakes in August 2022. Open circles: lakes with anoxic zone SS4 (orange) and SS6 (pink), closed circles: holomictic lakes SS3 (green) and SS5 (magenta).

### Sampling

Fieldwork was performed between 25^th^ July and 4^th^ August 2022. Oxygen saturation and temperature depth profiles were measured with a YSI 650 MDS multiprobe. Adult *Daphnia pulicaria* were collected from the water column with a 200 μm plankton net (diameter 25 cm). Additionally, we assessed the zooplankton depth distribution in lake SS4. For details see the supplementary material and Figure S1.

### Study Organism

Adult sampled *Daphnia pulicaria* were used to establish isofemale lineages, henceforth referred to as clones. *Daphnia* were cultured in SSS medium (Saebelfeld et al. 2017) at 14 °C under 24h dimmed light and fed with 1 mg C/L of *Tetradesmus obliquus* three times a week. The clones had been maintained in the lab for > 5 months (equal to approx. 7 generations) before measuring their respiration rate. For this study, we used four clones from each lake (Table S1).

### Whole genome sequencing (WGS)

*DNA extraction* - We attempted to genotype all clones measured in the respiration assay with WGS, but several clones from lakes SS3 and SS5 had gone extinct before DNA extraction. As replacement, we selected three substitute clones for lake SS3 and one for lake SS5 (see Table S1). Prior to DNA extraction, *Daphnia* were exposed to an antibiotics cocktail (50 mg/L ampicillin and 50 mg/L tetracycline dissolved in SSS-Medium) for 24 hours to remove the microbiome, and fed with Sephadex G-50 beads (5g/L) to clean the gut. Genomic DNA was extracted using the MasterPure^TM^ Complete DNA and RNA Purification kit according to the manufacturer’s instructions.

*Library preparation and sequencing* - DNA from 15 clones was used to prepare PCR-free paired-end sequencing libraries (Illumina DNA Prep PCR free) with an insert size of ∼300bp. Libraries were sequenced on the Illumina NovaSeq 6000 S4 Flowcell with 150bp paired-end-reads to an average of 6Gb per sample. Library preparation and sequencing were performed at the Competence Center for Genomic Analysis (CCGA) in Kiel, Germany.

### Bioinformatic analyses

Analyses involving the R platform were made with R version 4.4.1 (R Core Team 2023). All other bioinformatic analyses were completed within the High-Performance Computing infrastructure at ZEDAT, Freie Universität Berlin (Bennett et al. 2020).

*Short read mapping* - An initial quality check of paired-end fastq files was performed with FASTQC 0.11.9 (Andrews 2015). For adapter trimming we used TRIMGALORE 0.6.6 (Krueger et al. 2023) with the options “--length 35 and -q 20”. Paired-end reads were mapped to the *Daphnia pulex* reference genome (NCBI accession GCF_021134715.1) with BWA MEM 0.7.17 (Li and Durbin 2009) and default settings. From the mapped reads, duplicate reads and supplementary reads were removed with *markduplicates* from the PICARD Toolkit 2.26 (Broad Institute, 2024) and SAMTOOLS VIEW 1.10 (Danecek et al. 2021), respectively.

*Variant Call* - Variants were called with FREEBAYES 1.3.2 (Garrison and Marth 2012) with the following settings: --min-mapping-quality 40 --min-base-quality 24 --min-alternate-fraction 0.05 --min-alternate-count 5 -p 3 -g 10000. Variants were prefiltered with VCFFILTER FROM VCFLIB 1.0.3 (Garrison et al. 2022) with the settings “QUAL > 1”, “QUAL/AO > 10, “SAF > 0 & SAR > 0”, and “RPR > 1 & RPL > 1. Using the R package SEQARRAY 1.26.2 (Zheng et al. 2017), We selected only polymorphic, biallelic SNPs with MAF 0.15 and no missing data for further analysis. Finally, SNPs were selected with an average read depth between 10 and 100 reads per sample, resulting in the final SNP set of 368,820 SNPs.

The final SNP set was used as input to calculate pairwise genome-wide Identity-by-State (IBS) estimates. We adapted the formula of the *snpgdsIBS()* function in the R package SEQARRAY “mean(1-abs(x_i_-y_i_/2)” that estimates IBS for diploid genotypes to perform “mean(1-abs(x_i_-y_i_/3)”, where x and y represent the dosage of the reference allele of a pair of individuals at a given locus i as 0, 1, 2 and 3, divided by 3 (for three alleles instead of two). The obtained values are averaged across all SNP loci, resulting in a pairwise similarity matrix between clones. Based on the same input SNP set we also calculated a pairwise dissimilarity matrix with the formula “mean(abs(x_i_-y_i_ /3)” and used it for the hierarchical cluster analysis implemented in the R package SNPRELATE 1.38 (Zheng et al. 2012), with the functions *snpgdsHCluster()* and *snpgdsCutTree()*. Finally, we calculated the percentage of homozygosity across all identified SNPs for each member of the two genetic clusters.

*Functional analysis of SNPs -* Effects of the identified SNPs were predicted with SNPEFF 5.2 (Cingolani et al. 2012) with default settings. For further functional analysis we selected SNPs with a putatively moderate (missense variant, missense variant & splice region variant) or high impact (start or stop codon gained or lost, start lost & splice region variant, stop gained & splice region variant, stop lost & splice region variant, splice acceptor variant & intron variant, splice acceptor variant & splice donor variant & intron variant, splice donor variant & intron variant) and filtered the resulting gene list to contain only unique genes by removing duplicates caused by multiple SNPs on a gene in question.

*GO term enrichment -* We used the webtool EGGNOG-MAPPER 2.1.12 (http://eggnog-mapper.embl.de/; Cantalapiedra et al. 2021) with default settings for the GO annotation of the *Daphnia pulex* gene IDs (NCBI RefSeq assembly GCF_021134715.1_ASM2113471v1). The input file for EGGNOG-MAPPER were proteins of the *Daphnia pulex* genome GCF_021134715.1_ASM2113471v1_protein.faa.gz. To produce the final GO-annotated list of unique gene IDs, missing GO term entries were replaced by NAs for gene IDs that could not be mapped to any GO term. We used the list of unique genes that were affected by mutations with putatively *high* or *moderate* impact (see above). For comparison of a meaningful SNP subset between the two clusters, we produced two gene lists filtered to include genes in which the SNP was either

1. *homREFclusterB*: homozygous for the REF allele for clones of genetic cluster B and heterozygous or homozygous for the ALT allele in genetic cluster A (resulting in 3122 genes, final list with 1790 unique genes)
2. *homALTclusterB:* homozygous for the ALT allele in the two clones of genetic cluster B and heterozygous or homozygous for the REF allele in genetic cluster A (resulting in 322 genes, final list with 202 unique genes).

Both lists were used for enrichment against all 22,379 annotated genes in the *Daphnia pulex* ASM2113471v1 genome (NCBI RefSeq assembly GCF_021134715.1). Enrichment in GO terms was computed with the R package GOseq 1.56.0 (Young et al. 2010), with input gene set lists [1] and [2], the *Daphnia pulex* annotated gene list and the associated gene lengths. To calculate unbiased scores for GO term enrichment we used the default Wallenius non-central hypergeometric distribution. The resulting p-values for GO terms overrepresented in the input lists were adjusted by controlling for multiple comparisons (Benjamini and Hochberg 1995). Results were visualised with ggplot2 3.5.1 (Wickham 2016).

### Respirometer setup

Respiration was measured in a closed-chamber microplate respirometry system operated by the MicroResp^TM^ software (Loligo® Systems, Denmark) including a 200 μl 24-well plate, equipped with optical oxygen sensors and an SDR SensorDish® Reader (PreSens Precision Sensing GmbH, Germany). The plate was sealed with a PCR film and placed within a flow-through water bath for microplate (Loligo® Systems, Denmark) connected to a circulating refrigerated system (circulator: Thermo Scientific^TM^ HAAKE A10, thermostat: Thermo Scientific^TM^ HAAKE SC100) to keep a constant temperature at 14 ⁰C.

Oxygen diffusion rate at 14 ⁰C was determined prior to the assessment of respiration, by a 4h long measurement with oxygen saturated SSS medium, for correction of respiration rates (see below). The respiration rates were measured in five replicate runs. Within each experimental run, we used one replicate of each of the 16 clones, placing one sub-adult *Daphnia* per well. Additionally, four empty wells with an addition of a droplet of SSS medium from the cultures were used to account for background microbial respiration. To assess variation between runs, we included a reference clone present with four replicates in all five runs. The run effect was accounted for in the analysis using linear mixed models (see below).

Following each run, the body length of individual *Daphnia* was measured using an OLYMPUS SZX16 stereomicroscope and the OLYMPUS Stream Basic 2.5.2 imaging analysis software. Body length was measured from the top of the head (above the eye) to the frontal base of the apical spine.

### Statistical analysis of respiration rates

All analyses were performed in R (v. 4.3.0; R Core Team 2023). Raw data generated by the MicroResp^TM^ software were analyzed using the respR package (v. 2.3.1; Harianto et al. 2019). Background respiration rates were calculated separately per run as an average of the four control wells and were subtracted from the *Daphnia* respiration rates. An average of all wells from the diffusion measurement was used to further adjust the respiration rates. Respiration rates were adjusted by body mass using the length-weight ratio estimated from *Daphnia* from lake SS4 (for details see Supplementary methods). The resulting values are respiration rate per mg, henceforth referred to as Resp_adj_.

For the statistical analysis, we fitted general linear mixed effect models using the lmerTest (v. 3.1-3; Kuznetsova et al. 2017) and lme4 (v. 1.1-33; Bates et al. 2015) packages. For model L1 “Lake” was included as fixed effect, and “Run” as random effect. Model L2 included “Clone” as an additional fixed effect. Similarly, we fitted two models with the genetic clusters identified in this study as fixed effect instead of lake origin (model G1 and G2):

model L1: Resp_adj_ ∼ Lake + (1|Run)

model L2: Resp_adj_ ∼ Lake + Clone + (1|Run)

model G1: Resp_adj_ ∼ Genetic Cluster + (1|Run)

model G2: Resp_adj_ ∼ Genetic Cluster + Clone + (1|Run)

We compared all models and a null model (Resp_adj_ ∼ 1 + (1|Run)) using the Akaike Information (AIC) and Bayesian Information (BIC) criteria. Post-hoc tests corrected for multiple comparison with the Holm method were conducted with the multcomp package (v. 1.4-25; Hothorn et al. 2008).

For data visualization, we used the ggplot2 (v. 3.5.1; Wickham 2016) and patchwork (v. 1.2.0; (Pedersen 2024) packages.

## Results

### Two genomic clusters across four lakes

We performed whole genome sequencing for 15 clones of which 11 participated in the respiration assay (Table S1). Three clones from lake SS3 and two clones from lake SS5 used in this assay had expired prior to sequencing and were replaced by three and one clones for lakes SS3 and SS5, respectively. This was done with the intention to capture possible additional genomic variation of the lineages collected from the respective lakes. Comparing 15 clonal lineages from four lakes, we identified a final list of 368,820 high-confidence SNPs. Identity-by-state (IBS) analysis revealed two closely related genetic clusters (Figure 2A, clusters A and B). Cluster A comprised clones from all four lakes, while cluster B included only two clones from lake SS4. Pairwise dissimilarities (Fig. 2A) and similarities (Fig. 2B, supplementary Table S2) are calculated considering only the 368,820 identified SNPs. The highest pairwise dissimilarity between members of cluster B to members of cluster A was 9.6%, corresponding to *ca.* 0.28 % of the total number of bases across the genome, considering the size of the *Daphnia pulex* reference genome (about 133 Mb). Comparing the SNP set between the two clusters, we detected a nearly 4-fold higher homozygosity in the two clones of cluster B (Fig.S2).

**Figure 2.**
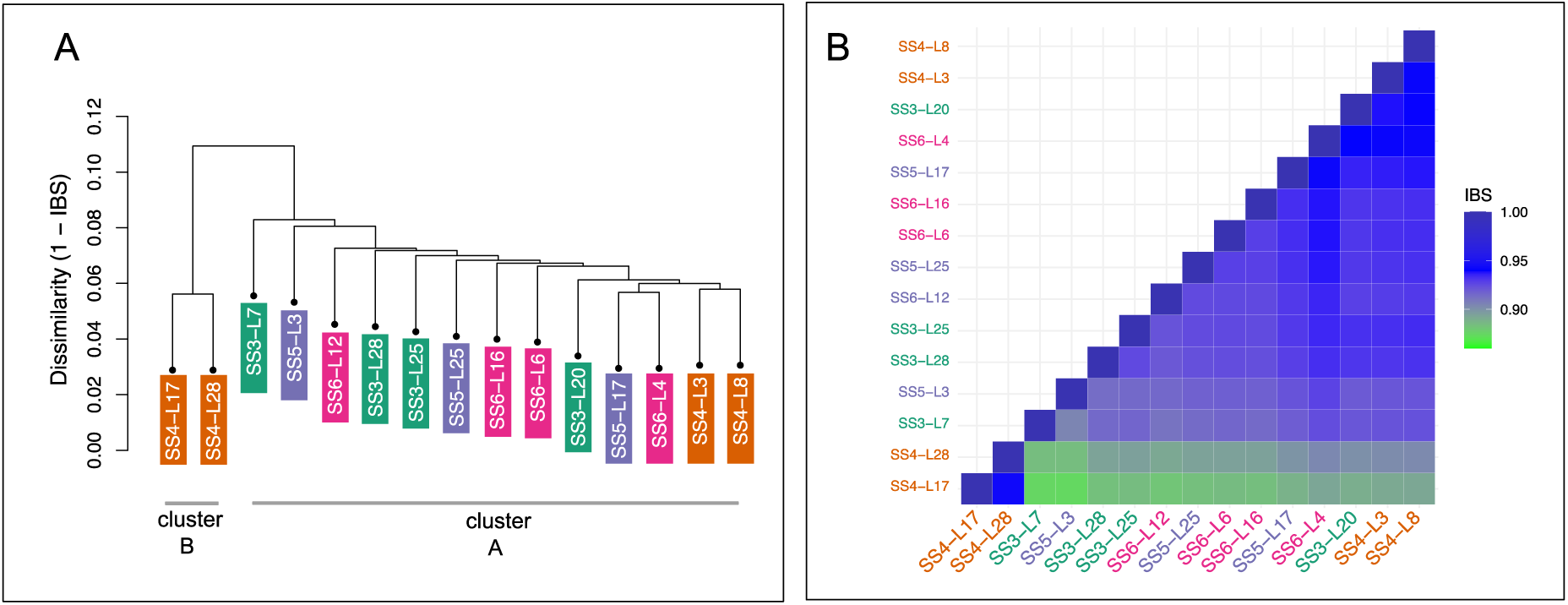
Clustering of clonal lineages using identity-by-state (IBS) analysis. A) Dendrogram representing the percentage of dissimilarity (1-IBS) between clonal lineages, based on 376,958 SNPs. B) Heatmap representing IBS in pairwise comparison between clonal lineages.

### Functional enrichment of SNPs in coding regions

To obtain a better understanding of genomic differences with possible functional impact between the two observed genetic clusters, we used the software SNPEFF to predict high (e.g. gained or lost stop codons) and moderate (e.g. missense mutations) effects of SNPs in coding regions. Focussing on loci homozygous for the reference allele only in the two clones of cluster B, we identified 189 loci with high and 2933 loci with moderate impact in the *homREFclusterB* list (Table S3). In the list *homALTclusterB* containing loci homozygous for the variant allele only in cluster B, a smaller number of high impact (7) and of moderate impact (315) loci were identified (Table S4). Because some of the genes contained multiple SNPs with high or moderate impact, we chose only unique entries for the GO term functional enrichment, with a total number of 1790 genes for gene list *homREFclusterB* and 202 genes for gene list *homALTclusterB*.

Functional analysis of the gene list *homREFclusterB* resulted in enrichment of 19 terms in the GO category biological process (BP), 7 terms in the category cellular component (CC) and 6 terms in the category molecular function (MF) (Fig. 3). Many of the enriched terms were more generally related to protein metabolism (e.g. proteolysis in BP and peptidase activity in MF), but also more specifically to nucleocytoplasmic transport (BP) and membrane-bounded organelle (CC), a term that refers to the nucleus and other organelles bounded by a membrane. The most highly enriched term, but also the one with the least number of genes was regulation of DNA methylation-dependent heterochromatin formation.

**Fig. 3.**
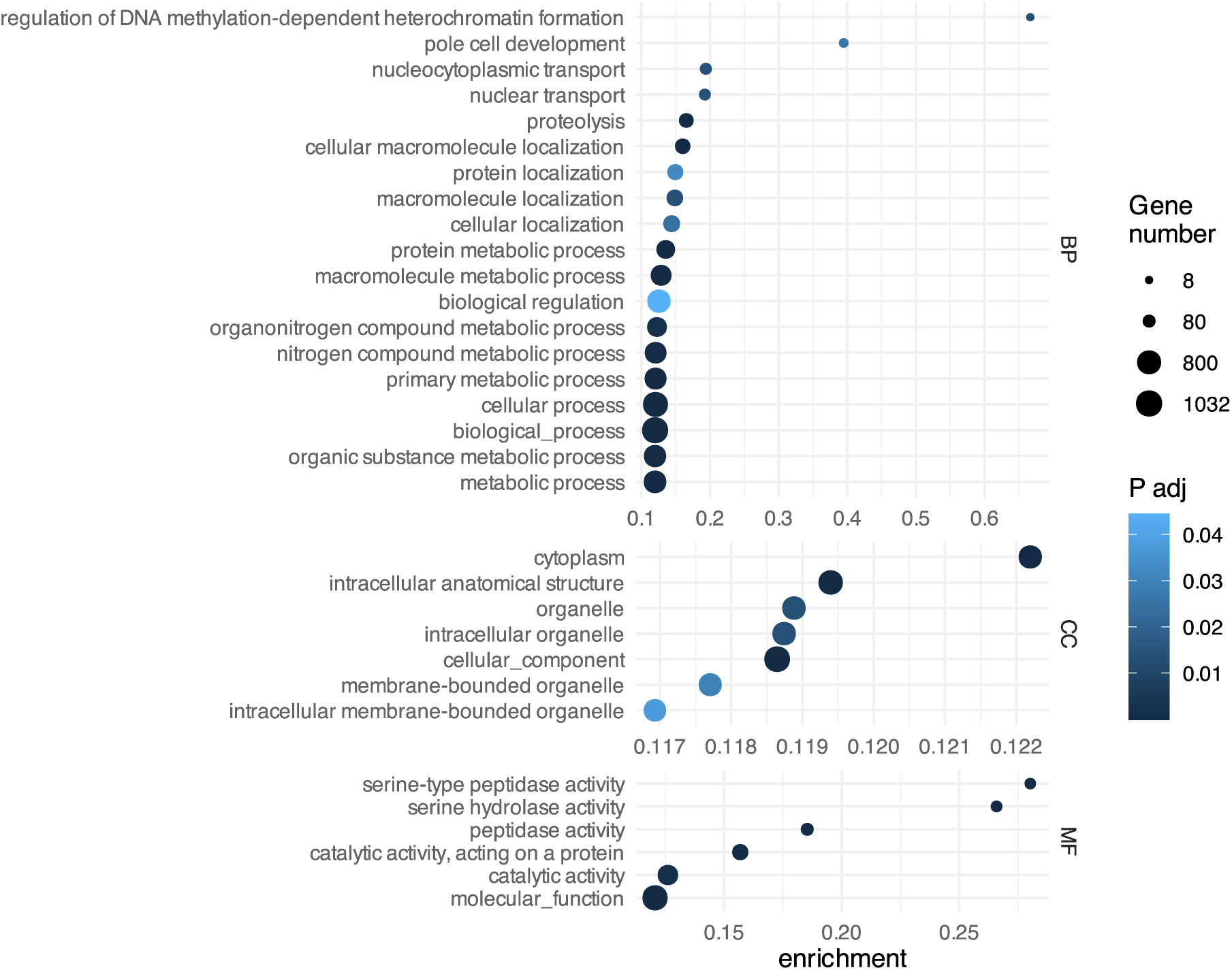
GO term enrichment of the SNPEFF-filtered gene list *homREFclusterB* with genes homozygous for the REF allele in genetic cluster B, and heterozygous or homozygous for the ALT allele in genetic cluster A. From top to bottom the three panels show GO categories BP (biological process), CC (cellular component) and MF (molecular function). The size of circles represents the number of genes in each category present in *homREFclusterB*. The enrichment score (x-axis) is the ratio between the number of genes in *homREFclusterB* and in the *Daphnia pulex* reference for each category. See methods for details.

In contrast, the gene list *homALTclusterB* yielded only a few enriched GO terms (Fig. S3), and similarly, most of the enriched terms were related to proteolysis (BP) and peptidase activity (MF).

### Respiration rates from four lakes

Two clones from lakes SS4 (genetic cluster B) had significantly lower respiration rates compared with clones from genetic cluster A (nine clonal lineages from all four lakes, Table 1a, Fig. 4a, and Table S5a). Respiration rates of five clones for which genomic data was not available (”Unknown” cluster) did not differ from those of cluster A (Fig 4a, Table S5a). The linear mixed model that included *genetic cluster* as fixed effect (G1) was selected by model comparison based on values for both AIC and BIC (Table 1c). As the genomic data did not include all clones participating in the assessment of respiration rates, we also show the results for model L1, which was the second best model according to the AIC and BIC values, and included *Lake* as fixed effect. The respiration rate of the clones from Lake SS4 was significantly lower in comparison with the other three lakes (Table 1a, Figure 4b, and Table S5b), while lakes SS6, SS3 and SS5 were similar to each other.

**Fig. 4.**
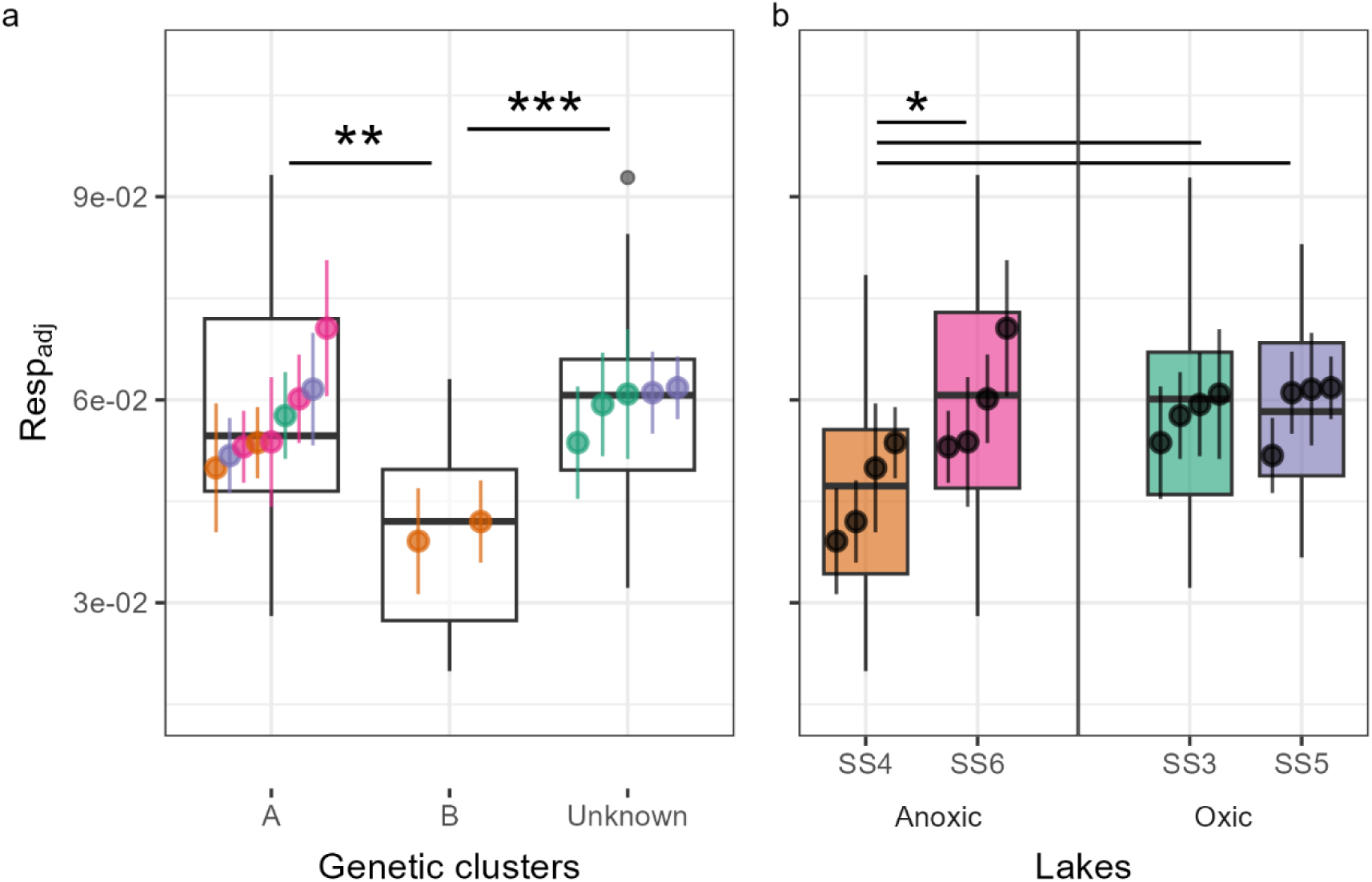
Respadj (dry weight-adjusted respiration rate in mg) of clones grouped by genetic cluster (a) and lake (b). **a**. Cluster A includes most clones from all lakes, while cluster B includes two clones from Lake SS4. The “Unknown” cluster consists of the clones from SS3 and SS5 for which genomic data was not available. **b**. Each lake includes four clones. Lakes SS4 and SS6 have an anoxic zone, while lakes SS3 and SS5 are holomictic. **Boxplots**: the center line is the median of each lake or cluster, box limits are the first and third quartiles, whiskers represent the 1.5 x interquartile range, grey points show outliers. **Filled circles**: mean of one clone across five experimental runs, error bars: standard error of the mean. Comparisons between groups (post-hoc test: Holm method) represented by horizontal bars. Statistical significance of p-value is shown by asterisks (* < 0.05, ** < 0.01, *** < 0.001). All three comparisons shown in b had the same significance level.

**Table 1.** **a.** Linear mixed model G1: Resp_adj_ ∼ gen. cluster + (1|Run) with genetic cluster as fixed effect and measurement run as random effect. Each cluster is compared with cluster A. The “Unknown” cluster contains the clones for which genomic data was not available. **b.** Linear mixed model L1: Resp_adj_ ∼ Lake + (1|Run) with lake as fixed effect and measurement run as random effect. Each lake is compared with Lake SS3. **c**. Model comparison using the Akaike Information Criterion (AIC) and Bayesian Information Criterion (BIC).

|  | Estimate | Std. Error | df | t value | Pr(>\|t\|) |
| --- | --- | --- | --- | --- | --- |
| *Intercept* | 0.057 | 0.004 | 6.147 | 13.595 | < 0.0001 |
| genetic cluster B | -0.016 | 0.005 | 74.972 | -3.462 | 0.0009 |
| genetic cluster "Unknown" | 0.003 | 0.003 | 75.042 | 0.786 | 0.4343 |

**b. Model L1:** $\text{Resp}_{\text{adj}} \sim \text{Lake} + (1 | \text{Run})$
|  | Estimate | Std. Error | df | t value | Pr(>\|t\|) |
| --- | --- | --- | --- | --- | --- |
| *Intercept* | 0.058 | 0.005 | 10.283 | 12.196 | < 0.0001 |
| Lake SS4 | -0.012 | 0.004 | 75.042 | -2.768 | 0.0071 |
| Lake SS5 | 0.001 | 0.004 | 75.042 | 0.207 | 0.8370 |
| Lake SS6 | 0.001 | 0.004 | 75.042 | 0.286 | 0.7755 |

**c. Model comparison**
|  | df | AIC | BIC |
| --- | --- | --- | --- |
| Null: $\text{Resp}_{\text{adj}} \sim 1 + (1 \vert \text{Run})$ | 3 | -432.433 | -425.286 |
| L1: $\text{Resp}_{\text{adj}} \sim \text{Lake} + (1 \vert \text{Run})$ | 6 | -438.431 | -424.139 |
| **G1: $\text{Resp}_{\text{adj}} \sim \text{gen. cluster} + (1 \vert \text{Run})$** | **5** | **-442.061** | **-430.15** |
| L2: $\text{Resp}_{\text{adj}} \sim \text{Lake} + \text{Clone} + (1 \vert \text{Run})$ | 18 | -427.35 | -384.474 |
| G2: $\text{Resp}_{\text{adj}} \sim \text{gen. cluster} + \text{Clone} + (1 \vert \text{Run})$ | 18 | -427.35 | -384.474 |

## Discussion

In this study we tested local adaptation of 16 triploid OP *Daphnia pulicaria* clones, from four lakes with two contrasting environments using respiration rate as a phenotypic trait combined with whole genome sequencing. We hypothesized the existence of local adaptation to the respective environments. We expected to find lake-specific genotypes that would have higher hypoxia tolerance in lakes SS4 and SS6 (with an anoxic zone and purple sulfur bacteria) than clones from the oxic lakes SS3 and SS5. Our findings confirmed that on average, clones from Lake SS4 had lower respiration rates than clones from the other three lakes but that this lower respiration rate was driven specifically by two clones belonging to a separate genetic cluster, indicating their local adaptation to environmental conditions in SS4.

### Two genetic clusters - evidence of local adaptation?

Previous studies in Greenland in other areas of West Greenland identified a high variation of asexual lineages in neighbouring ponds and lakes using allozyme markers and microsatellites (Weider et al. 1996; Haileselasie et al. 2016b). Contrary to their results we detected only two genetic clusters with a very high similarity to each other. Whole genome sequencing revealed the presence of two main genetic clusters of which the larger one (cluster A) contained 13 clones that had been sampled from all four lakes, while the smaller cluster B contained only two clones originating from lake SS4. SNP analysis showed a genome-wide difference between these two clusters of only *ca.* 0.3 %, however, respiration rates differed significantly between the two clusters, suggesting a genetic basis of this phenotypic trait. Detected in all four lakes of our study with different environmental regimes, clones of cluster A may be examples of a general purpose genotype (Lynch 1984), hypothesized to be a common characteristic of OPs. According to theory, since the whole genome is a single linkage group in asexual lineages, general purpose genotypes with higher fitness would be selected over a broader range of environments, realised by higher phenotypic plasticity. Field and experimental studies have provided evidence both for and against the general purpose genotype hypothesis (Vrijenhoek and Parker 2009). Lynch (1984) did not exclude the frequent appearance of specialist asexual genotypes but pointed out that these would be short-lived and become extinct as soon as the environmental niche that favoured them no longer exists. Other work (Haileselasie et al. 2016a) found support for the frozen niche variation model (Vrijenhoek 1984), where the repeated transition to asexuality from a sexual relative has frozen the genetic variation of each lineage, producing multiple clones with a specialized niche. Jose and Dufresne (2010) suggested that the presence of either specialists or generalists can be derived from clonal turnover: Higher clonal turnover would indicate specialist lineages and vice versa. It has been argued that patterns of local adaptation found in OP *Daphnia* populations. e.g. clonal genetic differentiation along a salinity gradient (Weider and Hebert 1987) can be achieved only by immigration of pre-adapted genotypes rather than local genetic differentiation (De Meester 1996a; De Meester et al. 2002). Tian et al. (2019) found evidence for molecular evolution via few high effect mutations in Japanese *Daphnia pulex* lineages. Our results show a very high similarity between clusters but similarly suggest that a group of novel mutations between the clusters could be functionally significant.

### Functional enrichment points to adaptation

Enrichment of cluster B genes with homozygous SNPs of high or moderate impact suggest a relationship of the involved genes in several functional processes. However, any potential impact on gene expression or translation must be interpreted with caution and requires future investigation, e.g. by comparing transcriptomic profiles of the two genetic clusters. Genes that were homozygous both for the reference and the variant allele in the individuals from cluster B, were mostly enriched in metabolic processes, in particular related to proteolysis and peptidase activity. This might suggest an evolution of alternative metabolic pathways utilised by representatives of the two genetic clusters to optimise digestion of available food items. Such a scenario could be similar to the findings of Tian et al. (2019) who detected significant genotype by food interactions in OP *Daphnia pulex* populations. Gene expression studies showed that different enzymatic pathways are utilised in the same *Daphnia* genotype when feeding on different diets (Koussoroplis et al. 2017; Schwarzenberger and Fink 2018). Previous studies have shown that purple sulfur bacteria (PSB) are consumed directly or indirectly by *Daphnia* (Jürgens et al. 1994; Massana et al. 1994). *Daphnia* feeding on anoxygenic photoautotrophic green sulfur bacteria was evidenced by biomarkers (Taipale et al. 2009). In our study, cluster B clones that also had a lower respiration rate compared with cluster A clones might be specifically adapted to exploit the PSB present in or near the oxycline of lake SS4 as a potential food source. In addition, enrichment analysis suggests the involvement of genes homozygous for the reference allele in epigenetic mechanisms, such as DNA methylation-dependent heterochromatin formation. Heterochromatin formation in *Daphnia magna* is closely related to histone modification and might play an important role in the epigenetic machinery of *Daphnia* (Robichaud et al. 2012). As with other enriched functional gene categories identified in our study, these might be possible candidates for local adaptation in the asexual populations of the study area and require closer examination in future research.

### Phenotypic and genotypic divergence match

While our genomic analyses did not identify functional enrichment in oxygen metabolism, the differences in respiration rates could be directly or indirectly related to an adaptation to hypoxia. Some of the physiological responses to chronic hypoxia in *Daphnia* include an elevation of hemoglobin production and higher oxygen affinity, but also a depression of metabolism that can lead to lower oxygen consumption (Wiggins and Frappell 2000; Paul et al. 2004). A central mechanism of response to hypoxia is the upregulation of hemoglobin production (Paul et al. 2004). It has been found that hypoxia-acclimated animals have a higher hemoglobin content as well as a lower respiration rate (reviewed in Zeis 2020). According to the above, the significantly lower respiration rates even at the normoxic conditions offered during the experiment of two *Daphnia* clones from Lake SS4 point to their higher hypoxia tolerance in comparison to clones from the other three lakes. Together with our observation of *Daphnia* being present across the entire water column including hypoxic and anoxic areas, we propose that at least some of the *Daphnia* can feed under hypoxic conditions for limited time periods.

A plausible explanation for the *Daphnia* migrating as far as the anoxic zone would be their exploitation of PSB as a food source. Previous sediment records have shown a correlation between the abundance of ephippia (a proxy of population size) and PSB abundance through time, suggesting a trophic link (Dane et al. 2020). It is known that *Daphnia* can graze on PSB in the field (Jürgens et al. 1994; Massana et al. 1994), however, the extent to which they might depend on such a food source is still unknown. Notably, in oligotrophic lakes such as those studied here, the abundance of other food sources might not suffice during the growing season, driving the adaptation to hypoxia which allows exploitation of PSB as an important food source for some of the clones.

We expected to find a similar adaptation to hypoxia tolerance in *Daphnia* of lakes SS4 and SS6. However, the respiration rates of SS6 clones were similar to those of SS3 and SS5 reflecting their membership to genetic cluster A. Since oxygen in SS6 did not drop to complete anoxia as it does in SS4, the conditions might not be ideal for the formation of extensive PSB plates and hence cluster B might not be adaptive in lake SS6. Additionally, SS4 with its higher heterogeneity, specifically regarding oxygen conditions, would allow for niche compartmentalization, and hence for members of both genetic clusters to coexist.

## Conclusion

Our study is focused on only four lakes with a limited number of clonal lineages. It is possible that additional, undetected genotypes exist in these lakes. However, our results are in agreement with the low variation found in the eggbank of lake SS4, where only two genotypes per temporal subpopulation were detected across hundreds of years (Dane et al., 2020). Although we cannot exclude the possible presence of cluster B not only in SS4 but also in the other lakes of this study, our data at least approximate relative clonal variation among the four lakes. Additionally, the use of whole genome sequencing has offered better resolution of the clonal clusters and unique insights into their possible functional differences.

In conclusion, we observed local adaptation of OP *Daphnia* to the distinct environmental conditions of lake SS4. Lower respiration rates, intimately related to metabolism, may reflect their frequent presence at or near the oxycline, which in this oligotrophic system can be interpreted as utilizing PSB as a direct or indirect food source. In addition, our whole genome data indicated molecular evolution in the studied OP *Daphnia* lineages but also raise questions on the mechanistics of phenotypic adaptation in asexual clones, including the possibility of epigenetic regulation of transcription.

## Acknowledgements

This work was supported by the DFG Research Infrastructure NGS_CC (project 407495230) as part of the Next Generation Sequencing Competence Network (project 423957469). NGS analyses were carried out at the Competence Centre for Genomic Analysis (Kiel). We thank James Shilland and Dörthe Becker for their invaluable discussions and their assistance in the field. We are grateful to the Government of Greenland for granting Prior Informed Consent for utilization of Greenland genetic resources (non-exclusive licence no. G22-076) that permits collection and basic research on genetic resources.

## Funding

This research was funded by the Deutsche Forschungsgemeinschaft (DFG, German Research Foundation) – Project number 461099895

## Author contributions

**AKP** Conceptualization; Formal analysis; Investigation; Data Curation; Visualization; Methodology; Writing - Original Draft; Writing - Review & Editing

**JJH** Formal analysis; Investigation; Writing - Review & Editing

**NJA** Conceptualization; Resources; Supervision; Writing - Original Draft; Writing - Review & Editing

**SF** Investigation; Resources; Writing - Review & Editing

**DF** Conceptualization; Formal analysis; Investigation; Resources; Data Curation; Visualization; Methodology; Supervision; Project administration; Funding acquisition; Writing - Original Draft; Writing - Review & Editing

## Notes

### Competing Interest Statement

The authors have declared no competing interest.

