## Supplementary material for "Phenotypic divergence associated with genomic changes suggest local adaptation in obligate asexuals"

### Methods

#### Study lake depth profiles

The recorded oxygen saturation and temperature along the water column across the four lakes on the day of sampling are shown in Figure 1. Close to 100% oxygen saturation was measured in all four lakes at 0 m depth. Oxygen saturation in Lake SS4 reached 0% at 13 m depth and the water column remained anoxic down to the sediment at 22 m water depth. Lake SS6 did not become completely anoxic but oxygen dropped to 15% at its deepest point of 13 m. In contrast, Lakes SS3 and SS5 retained oxic conditions throughout the water column with a minimum of 70% at 19 m. Temperature ranged from 14-15 $^{o}$C at the water surface to 3-4 $^{o}$C near the sediment and was similar across lakes.

#### Zooplankton depth distribution

We assessed the zooplankton distribution across the water column in lake SS4 with a 5 L Van-Dorn Sampler (height 41.5 cm, diameter: 13 cm), sampling every 1 m between 0 and 22 m (lake bottom). For a finer resolution throughout the oxycline, we sampled every 0.5 m between 10 and 14 m water depth. The PSB population in SS4 and SS6 using the Van-Dorn water sampler were not possible to locate during sampling. Samples were preserved in ethanol for later identification. Zooplankton species were identified according to (Røen 1962; Voigt and Koste 1978; Einsle 1993; Flössner 2000; Thorp and Rogers 2016).

#### Length-Weight transformation formula

Dry weight  was estimated from length measurements by linear regression of log-transformed dry weight and length of 104 total *Daphnia* originating from clones of lake SS4 and ranging from 792 to 2783 µm in size. We measured body length of live *Daphnia* from the top of the head above the eye to the ventral base of the apical spine using a dissecting scope (ZEISS, STEMI 508) at a magnification of 25x and the image analysis software MikroLive v.5 (https://www.mikroskopie.de/). To measure dry weight we placed *Daphnia* in aluminium cups of 3 mm diameter (one individual per cup). Prior to this, the cups were dried at 60 $^{0}$C for 17 hours and transferred into an exicator connected to a vacuum pump at 200 bar for an hour to achieve room temperature before weighing at a high-precision scale (Sartorius Cubis MSE2.7S-000-DM Micro Balance). Each cup was weighed three times to account for measuring error and the average was used for further analysis. Individual *Daphnia* were placed into the pre-weighed tins using fine forceps and weighed after drying as explained above. We calculated the relationship of dry weight to length by fitting a linear model of log(Dry Weight) ~ log(Length) using the lme4 package (v. 1.1-33; Bates et al. 2015). in R (v. 4.3.0; R Core Team 2023). The resulting formula (Dry Weight = 9.015362 * Length$^{2.86448}$, d.f. = 102, $r^2$ = 0.8259, P <0.001) was further used for assessing dry weight from length measurements in this study.

### Results

**Zooplankton is present in anoxic zone in SS4**

In SS4, six zooplankton species were identified. Of these, three species belong to the Cladocera: *Daphnia pulicaria* Forbes, 1893, *Chydorus sphaericus* (O.F. Müller*,* 1776) and *Polyphemus pediculus* (Linnaeus, 1761). The other three were a calanoid (*Leptodiaptomus minutus* (Lilljeborg, 1889)), a cyclopoid copepod (*Cyclops abyssorum* Sars, G.O. 1863) and one monogonont rotifer (*Brachionus cf. diversicornis* (Daday, 1883)). Most of these occurred only in small numbers and closer to the water surface, with the exception of the two copepod species and *Daphnia pulicaria*.  The distribution of these more abundant species is shown in Figure S1. Calanoid and cyclopoid copepods were more abundant than *Daphnia*, but all three species were distributed across the entire water column, including the anoxic zone.

Supplementary Figures

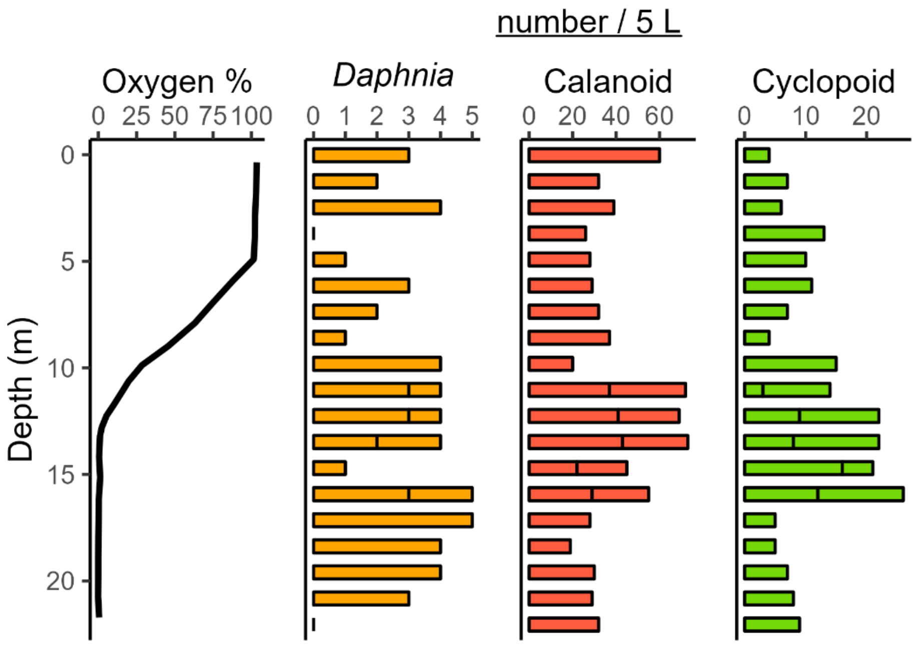


Fig. S1 Zooplankton total counts per 5 L for each meter from the surface (0m) to the bottom (22m) of lake SS4. Between 10 and 16 m the samples were taken every 0.5 m. The count per sample is shown by the vertical lines on the bars. Both adults and juveniles are included in the count.

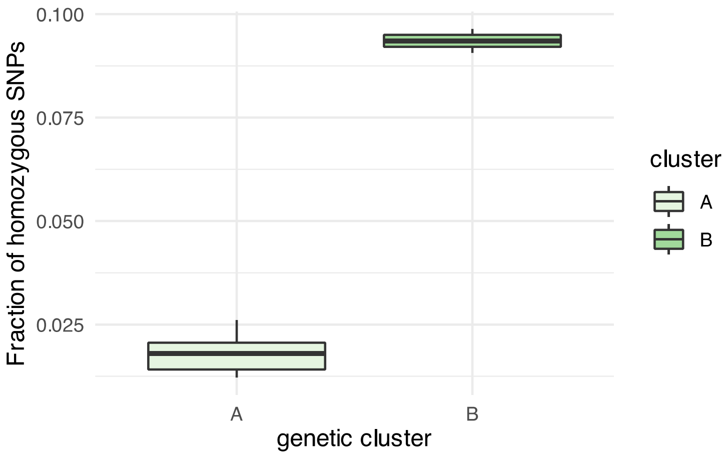


Figure S2. Fraction of homozygous SNPs of the total number of high confidence SNPs between cluster A and B

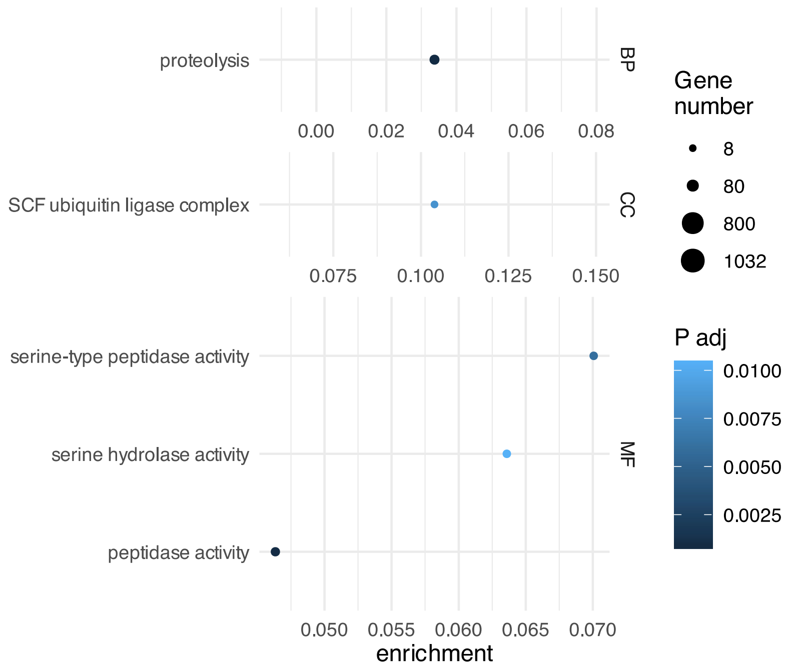


Figure S3. GO term enrichment of the SNPEFF-filtered gene list *homALTclusterB* with genes homozygous for the ALT allele in genetic cluster B, and heterozygous or homozygous for the REF allele in genetic cluster A. From top to bottom the three panels show GO categories BP (biological process), CC (cellular component) and MF (molecular function). The size of circles represents the number of genes in each category present in *homALTclusterB*. The enrichment score (x-axis) is the ratio between the number of genes in *homREFclusterB* and in the *Daphnia pulex* reference for each category. See methods for details.

### Supplementary tables

Table S1. Clones per lake used for the respiration assay and for whole genome sequencing.

| Clone | Lake | Respiration rate measurement | DNA genomic sequence available |
|---|---|---|---|
| SS3-L7 | SS3 | yes | yes |
| SS3-L15 | SS3 | yes | no |
| SS3-L26 | SS3 | yes | no |
| SS3-L30 | SS3 | yes | no |
| SS3-L20 | SS3 | no | yes |
| SS3-L25 | SS3 | no | yes |
| SS3-L20 | SS3 | no | yes |
| SS4-L3 | SS4 | yes | yes |
| SS4-L8 | SS4 | yes | yes |
| SS4-L17 | SS4 | yes | yes |
| SS4-L28 | SS4 | yes | yes |
| SS5-L3 | SS5 | yes | yes |
| SS5-L12 | SS5 | yes | no |
| SS5-L17 | SS5 | yes | yes |
| SS5-L29 | SS5 | yes | no |
| SS5-L25 | SS5 | no | yes |
| SS6-L4 | SS6 | yes | yes |
| SS6-L6 | SS6 | yes | yes |
| SS6-L12 | SS6 | yes | yes |
| SS6-L16 | SS6 | yes | yes |

Table S2. IBS Similarity matrix of pairwise similarities between the clones used in this study, condiering 368,820 high-confidence SNPs. For details see Methods in the main manuscript.

| | **SS4.L17** | **SS4.L28** | **SS3.L7** | **SS5.L3** | **SS3.L28** | **SS3.L25** | **SS6.L12** | **SS5.L25** | **SS6.L6** | **SS6.L16** | **SS5.L17** | **SS6.L4** | **SS3.L20** | **SS4.L3** | **SS4.L8** |
|---|---|---|---|---|---|---|---|---|---|---|---|---|---|---|---|
| **SS4.L17** | 1 | 0.944 | 0.875 | 0.875 | 0.884 | 0.885 | 0.882 | 0.884 | 0.885 | 0.884 | 0.889 | 0.893 | 0.890 | 0.892 | 0.893 |
| **SS4.L28** | 0.944 | 1 | 0.885 | 0.884 | 0.894 | 0.895 | 0.892 | 0.895 | 0.895 | 0.895 | 0.900 | 0.904 | 0.901 | 0.903 | 0.903 |
| **SS3.L7** | 0.875 | 0.885 | 1 | 0.905 | 0.915 | 0.917 | 0.912 | 0.915 | 0.916 | 0.915 | 0.919 | 0.924 | 0.922 | 0.923 | 0.923 |
| **SS5.L3** | 0.875 | 0.884 | 0.905 | 1 | 0.914 | 0.915 | 0.916 | 0.918 | 0.919 | 0.918 | 0.923 | 0.927 | 0.920 | 0.922 | 0.922 |
| **SS3.L28** | 0.884 | 0.894 | 0.915 | 0.914 | 1 | 0.926 | 0.921 | 0.924 | 0.924 | 0.924 | 0.929 | 0.933 | 0.931 | 0.931 | 0.932 |
| **SS3.L25** | 0.885 | 0.895 | 0.917 | 0.915 | 0.926 | 1 | 0.923 | 0.925 | 0.926 | 0.925 | 0.930 | 0.935 | 0.933 | 0.933 | 0.934 |
| **SS6.L12** | 0.882 | 0.892 | 0.912 | 0.916 | 0.921 | 0.923 | 1 | 0.925 | 0.926 | 0.926 | 0.930 | 0.935 | 0.928 | 0.929 | 0.930 |
| **SS5.L25** | 0.884 | 0.895 | 0.915 | 0.918 | 0.924 | 0.925 | 0.925 | 1 | 0.928 | 0.928 | 0.933 | 0.938 | 0.931 | 0.932 | 0.933 |
| **SS6.L6** | 0.885 | 0.895 | 0.916 | 0.919 | 0.924 | 0.926 | 0.926 | 0.928 | 1 | 0.929 | 0.933 | 0.938 | 0.931 | 0.933 | 0.933 |
| **SS6.L16** | 0.884 | 0.895 | 0.915 | 0.918 | 0.924 | 0.925 | 0.926 | 0.928 | 0.929 | 1 | 0.933 | 0.938 | 0.931 | 0.932 | 0.933 |
| **SS5.L17** | 0.889 | 0.900 | 0.919 | 0.923 | 0.929 | 0.930 | 0.930 | 0.933 | 0.933 | 0.933 | 1 | 0.943 | 0.936 | 0.937 | 0.938 |
| **SS6.L4** | 0.893 | 0.904 | 0.924 | 0.927 | 0.933 | 0.935 | 0.935 | 0.938 | 0.938 | 0.938 | 0.943 | 1 | 0.941 | 0.942 | 0.943 |
| **SS3.L20** | 0.890 | 0.901 | 0.922 | 0.920 | 0.931 | 0.933 | 0.928 | 0.931 | 0.931 | 0.931 | 0.936 | 0.941 | 1 | 0.939 | 0.940 |
| **SS4.L3** | 0.892 | 0.903 | 0.923 | 0.922 | 0.931 | 0.933 | 0.929 | 0.932 | 0.933 | 0.932 | 0.937 | 0.942 | 0.939 | 1 | 0.942 |
| **SS4.L8** | 0.893 | 0.903 | 0.923 | 0.922 | 0.932 | 0.934 | 0.930 | 0.933 | 0.933 | 0.933 | 0.938 | 0.943 | 0.940 | 0.942 | 1 |

*Tables S3 and S4 are provided in Excell format*

Table S5. Holm method post-hoc tests between a. genetic clusters and b. lakes. Significance codes: * < 0.05, ** < 0.01, *** < 0.001

a. Model G1: mass_adj_rates ~ gen_group + (1 | Run)

| **Comparison** | **Estimate** | **Std. Error** | **z value** | **Pr(>\|z\|)** |
|---|---|---|---|---|
| Cluster B - A | -0.016 | 0.005 | -3.462 | **0.0011 **** |
| Cluster unknown - A | 0.003 | 0.003 | 0.786 | 0.4318 |
| Cluster unknown - B | 0.019 | 0.005 | 3.757 | **0.0005 ***** |

b. Model L1: mass_adj_rates ~ Lake + (1 | Run)

| **Comparison** | **Estimate** | **Std. Error** | **z value** | **Pr(>\|z\|)** |
|---|---|---|---|---|
| SS4 - SS3 | -0.012 | 0.004 | -2.768 | **0.0225 *** |
| SS5 - SS3 | 0.001 | 0.004 | 0.207 | 1.0000 |
| SS6 - SS3 | 0.001 | 0.004 | 0.286 | 1.0000 |
| SS5 - SS4 | 0.013 | 0.004 | 2.979 | **0.0145 *** |
| SS6 - SS4 | 0.013 | 0.004 | 3.059 | **0.0133 *** |
| SS6 - SS5 | 0.000 | 0.004 | 0.080 | 1.0000 |
